# Coordinate genomic association of transcription factors controlled by an imported quorum sensing peptide in *Cryptococcus neoformans*

**DOI:** 10.1101/2020.03.30.016329

**Authors:** Diana K. Summers, Daniela S. Perry, Beiduo Rao, Hiten D. Madhani

## Abstract

Qsp1 is a secreted quorum sensing peptide required for virulence of the fungal meningitis pathogen *Cryptococcus neoformans.* Qsp1 functions to control cell wall integrity in vegetatively growing cells and also functions in mating. Rather than acting on a cell surface receptor, Qsp1 is imported to act intracellularly via the predicted oligopeptide transporter Opt1. Here, we identify a transcription factor network as a target of Qsp1. Using whole-genome chromatin immunoprecipitation, we find Qsp1 controls the genomic associations of three transcription factors to genes whose outputs are regulated by Qsp1. One of these transcription factors, Cqs2, is also required for the action of Qsp1 during mating, indicating that it might be a shared proximal target of Qsp1. Consistent with this hypothesis, deletion of *CQS2* impacts the binding of other network transcription factors specifically to Qsp1-regulated genes. These genetic and genomic studies illuminate mechanisms by which an imported peptide acts to modulate eukaryotic gene expression.

**AUTHOR SUMMARY:** For many fungal pathogens, the ability to adapt to changing and diverse environments forms the basis for their ability to infect and survive inside macrophages and other niches in the human body, and these changes are accomplished by transcription factors. Many pathogenic microbes coordinate their gene expression as a function of cell density in a process known as quorum sensing. Here, in the human fungal meningitis pathogen *Cryptococcus* neoformans, we find that an imported eukaryotic quorum sensing peptide that is important for virulence, Qsp1, controls the binding of three different transcription factors to promoters, thereby modulating the expression of Qsp1-regulated genes. This discovery reveals the mechanism for how an imported peptide affects gene expression.

## INTRODUCTION

The opportunistic basidiomycete yeast *Cryptococcus neoformans* is the most common cause of fungal meningitis, causing over 200,000 deaths annually (1). The unique features of this organism that drive its virulence are incompletely understood. In many bacterial pathogens, quorum sensing plays a key role in the regulation of group behaviors and virulence (2, 3). In previous work, we described a peptide-based quorum sensing system in *Cryptococcus neoformans*, the first described in a eukaryote (4). This system is mediated by an 11 residue peptide dubbed Qsp1, first purified because it complements a low-density phenotype produced by *C. neoformans* lacking a transcriptional co-repressor, Tup1 (5). Our analysis revealed that Qsp1 is secreted as a pro-peptide that is matured extracellularly by the cell wall-associated serine protease, Pqp1 into a biologically active form (4). The action of Qsp1 requires an oligopeptide transporter, Opt1 (4). As cytosolic expression of the mature form of Qsp1 complements the *qsp1*Δ knockout phenotype (a dry colony phenotype), we infer that Qsp1 acts intracellularly after import (4).

In this prior work, we demonstrated that a WOPR domain transcription factor, Liv3, which is related to key regulatory proteins *C. albicans* Wor1 and *H. capsulatum* Ryp1, acts downstream of Qsp1 (4,6–10). Like cells lacking other components of the Qsp1 system, cells lacking Liv3 display a rough colony morphology phenotype. Cells that lack Qsp1 or Liv3 are also attenuated for virulence in an intranasal mouse model of infection (4, 11). Others have discovered that Qsp1 also regulates unisexual and bisexual mating in *C. neoformans* as well as mating-induced transcription (12). This function also requires Opt1 and a previously uncharacterized transcription factor, Cqs2, which has also been called Zfc3 (12, 13). The relationships between the roles of Qsp1 in colony morphology, virulence, and mating are not well-understood.

In this paper, we demonstrate that mutants of two transcription factors in addition to Liv3 display a rough colony phenotype when deleted, Nrg1 and Cqs2. By performing a series of transcriptomics experiments, we show that these transcription factors and Qsp1 regulate a common set of target genes. Whole-genome chromatin immunoprecipitation demonstrates that these transcription factors generally bind together to a common set of target genes, forming a highly connected transcription factor network. Significantly, the presence of Qsp1 impacts the binding of all three transcription factors to a subset of target genes which are highly enriched for genes whose expression is controlled by Qsp1. Cqs2 is particularly sensitive to the presence of Qsp1 for its genomic binding. Cqs2 is strongly required for the binding of Nrg1 and Liv3 to Qsp1-regulated genes, suggesting it may be an upstream factor in the pathways. Furthermore, while Qsp1 seems to negatively regulate protein levels of Nrg1 and Liv3, the association of these factors with promoters is still greatly decreased in the *qsp1*Δ mutant. These experiments illuminate the mechanism by an imported quorum-sensing peptide impacts gene expression.

## RESULTS

### Phenotypic identification of predicted transcription factors that act downstream of Qsp1

Wild-type yeast form glossy colonies, whereas cells lacking the *QSP1* gene (*qsp1*Δ) exhibit a wrinkled colony morphology phenotype at either 25°C or 30°C (4). We previously published that the transcription factor Liv3 mediates a large portion of the Qsp1 response in rich media, and that a *liv3*Δ knockout strain forms dry, wrinkled colonies at 30°C. We hypothesized that the deletion of genes encoding factors involved in the response to Qsp1 signaling would also exhibit a wrinkled colony morphology. Therefore, we screened strains in a *C. neoformans* knockout collection generated in our laboratory for genetic candidates. We discovered two additional strains that exhibited a *qsp1*Δ-like colony morphology that is also temperature-dependent, corresponding to genes encoding the transcription factors Cqs2 and Nrg1. Cqs2 was recently reported as a regulator of the Qsp1 response for unisexual filamentation (12). Nrg1 is a transcriptional regulator that plays a role in several cellular processes, including carbohydrate acquisition, metabolism, and virulence (14).

In contrast to the *qsp1*Δ mutant, each transcription factor deletion strain exhibits this phenotype at a more restricted range of temperatures (Figure 1A). Colonies formed by *nrg1*Δ cells display their strongest phenotype at room temperature, and *liv3*Δ and *cqs2*Δ colonies show their strongest phenotype at 30°C. This dry and wrinkled colony morphology is not caused by an inability of these transcription factor deletion strains to synthesize Qsp1 peptide, as they are still able to secrete wild-type levels of Qsp1 peptide (Figure 1B). Additionally, each transcription factor deletion strain is able to complement a *qsp1*Δ strain when patched nearby on a plate, due to Qsp1 peptide diffusing through the agar (Figure 1C). In contrast, the colony morphology phenotype of the transcription factor deletion strains could not be complemented by the peptide produced by a wild-type strain (Figure 1C). Therefore, while each transcription factor knockout strain is able to produce Qsp1 peptide, none are complemented by the peptide. This supports the idea that all three of these transcription factors act downstream of Qsp1 production to promote wild type colony morphology.

**Figure 1.**
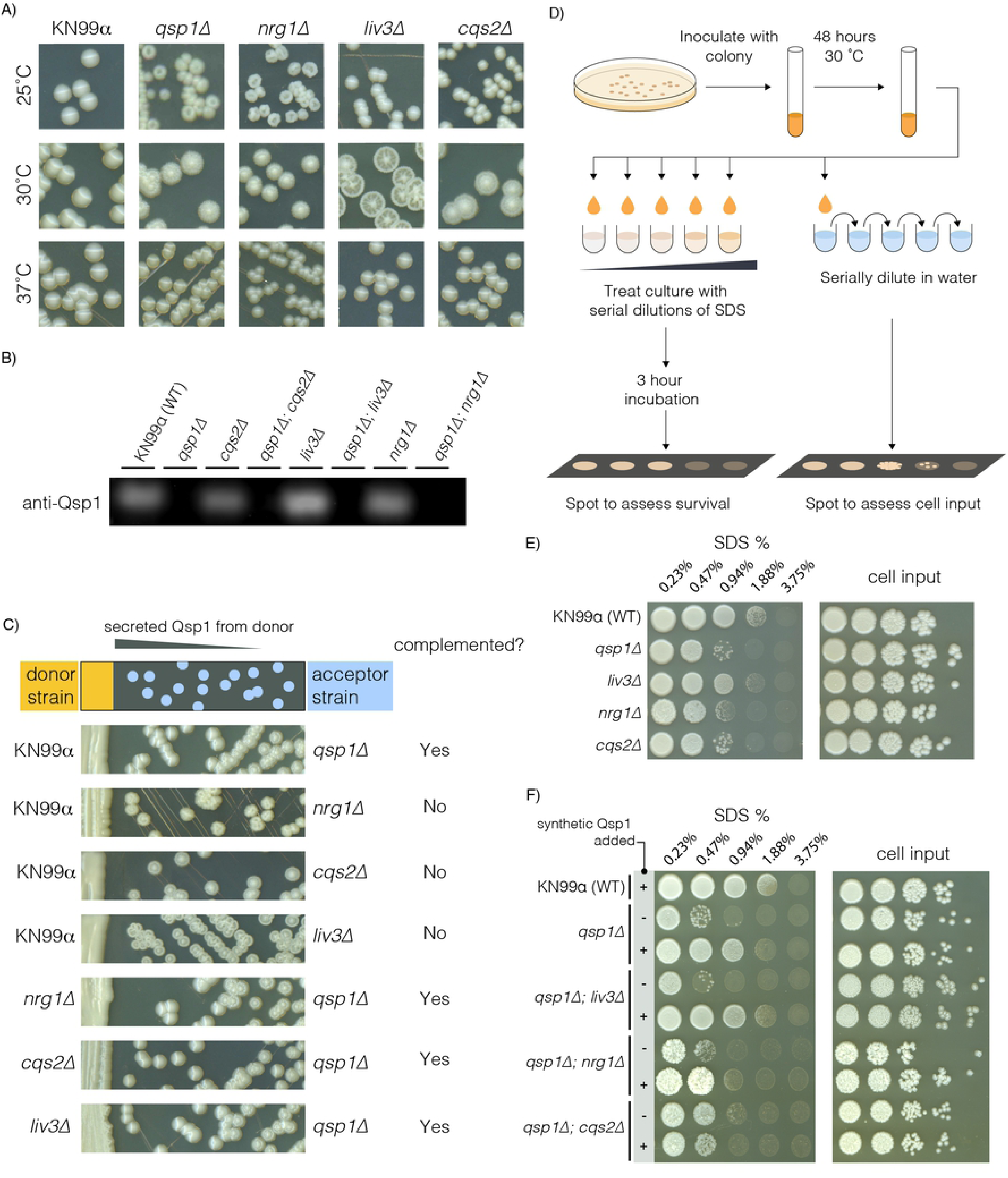
*LIV3*, *NRG1*, and *CQS2* act downstream of *QSP1*. A) Colony morphology of knockout strains streaked on YPAD agar at 25, 30, or 37°C. B) Anti-Qsp1 immunoblot of supernatant harvested from overnight cultures in YPAD. C) A Qsp1-secreting donor strain is patched on the left, and colonies from a wrinkled acceptor strain streaked to the right are tested for its ability to be complemented by Qsp1 from the donor strain. D) Schematic for how strains were tested for their ability to survive different concentrations of the cell wall stressor SDS. E) Single mutants for each of the genes shown were tested for their ability to survive increasing concentrations of the cell wall stressor SDS. Water dilutions are shown to the right. Plates were allowed to grow up at room temperature. F) Each genotype shown was tested for their ability to survive increasing concentrations of the cell wall stressor SDS. 1 uM synthetic Qsp1 peptide was added to the indicated cultures (+) from the time of inoculation, or not (-). Water dilutions of each culture are shown to the right as a measure of cell input.

Saturated cultures of the *qsp1*Δ mutant are sensitive to the cell wall stressor SDS, a phenotype that can be rescued by prior growth of the cells in the presence of synthetic Qsp1 peptide (4). To determine whether these three transcription factors could be involved in Qsp1 signaling in this context, we tested the sensitivity of the corresponding deletion mutants to SDS. We grew each strain to saturation in rich media, then incubated the cells in different concentrations of SDS (Figure 1D). The cells were then plated on YPAD agar to assay for viability following SDS treatment. The *liv3*Δ strain is not sensitive to SDS treatment, but *nrg1*Δ and *cqs2*Δ strains display sensitivity (Figure 1E). Thus, Nrg1 and Cqs2 function to promote resistance to cell wall stress, while Liv3 is dispensable for this phenotype.

To test whether Nrg1, Cqs2, or Liv3 were downstream of Qsp1, we created double knockouts of each transcription factor gene and *QSP1* and grew these strains with or without an excess of synthetic Qsp1 peptide for 48 hours (Figure 1F). The *qsp1Δnrg1*Δ double mutant is only modestly complemented, and the *qsp1Δcqs2*Δ double mutant is completely unable to respond to peptide. The SDS sensitivity of the *qsp1Δliv3*Δ mutant could be rescued by prior growth in synthetic Qsp1, indicating that Liv3 is not required the ability of Qsp1 peptide to promote SDS resistance (as expected from the lack of SDS sensitivity in the *liv3*Δ strain). These data are consistent with a model in which Cqs2 and Nrg1 function downstream of Qsp1 to promote resistance to a cell wall stress.

### RNA-seq analysis reveals shared roles for Qsp1 and the three transcription factors

In previous work, we found that loss of Liv3 significantly impacts the response of cells to Qsp1 (4). These experiments were performed at a single time point in rich media. To test more broadly media and culture density conditions for subsequent analysis, we collected RNA from either wild-type or *qsp1*Δ mutant cultures grown in either rich media (YPAD) or minimal media (YNB) at an optical density at 600 nm (OD_600_) of 1, 5 and 10 (Figure 2A). We then performed RNA-seq analysis to identify differentially expressed genes. Over 400 genes were significantly affected by the loss of the *QSP1* gene in minimal media at an OD_600_ of 1 (OD1) or OD_600_ of 5 (OD5), more genes than in rich media at any culture density (Figure 2B). Therefore, we chose to proceed with OD1 and OD5 conditions in minimal media for the subsequent experiment.

**Figure 2.**
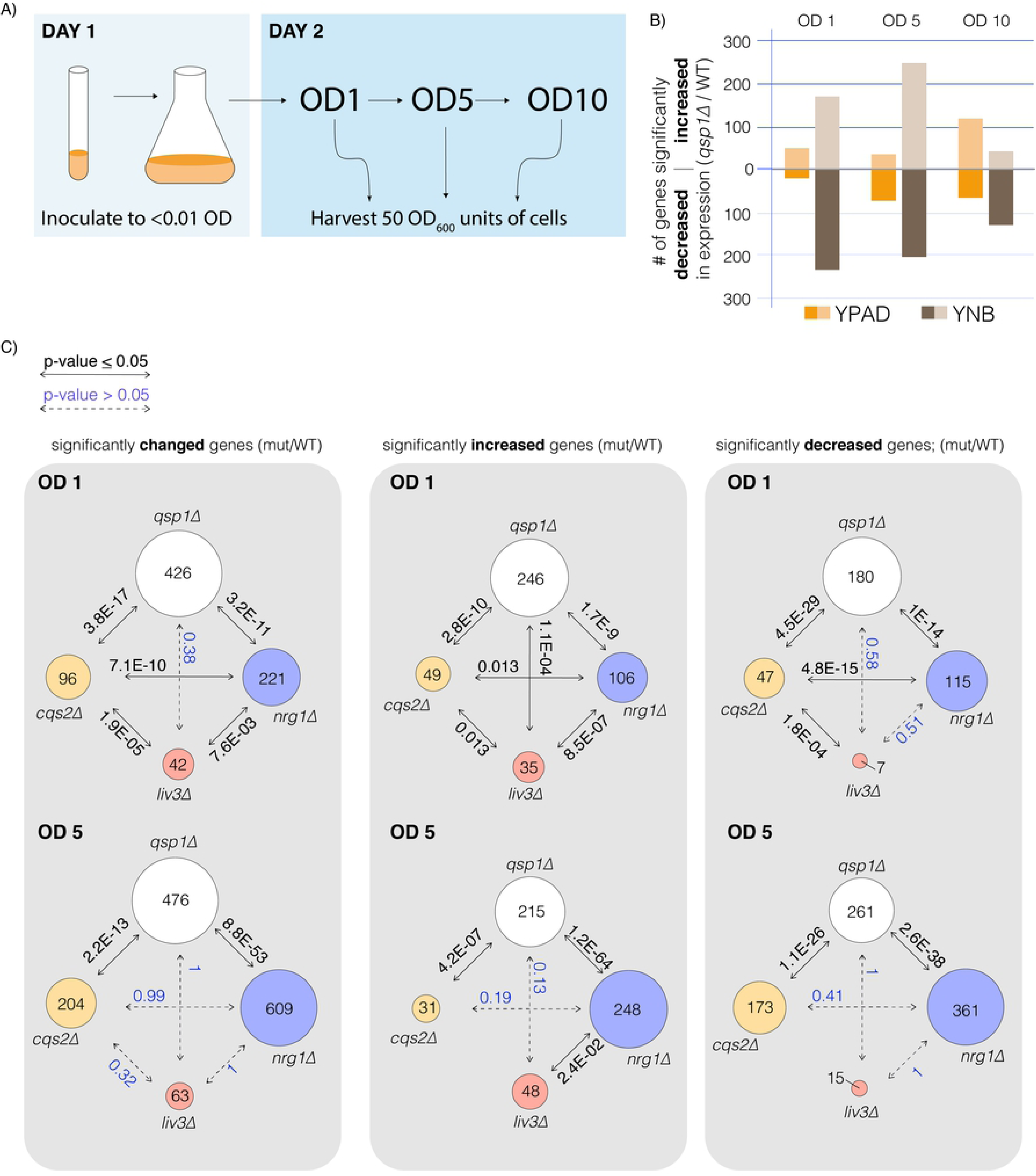
Cqs2, Nrg1, and Liv3 are part of a transcription factor network that shares targets with Qsp1. A) Schematic of how cultures were grown and harvested for RNA-seq. B) Number of significantly changed genes in *qsp1*Δ vs. wild type as determined by DE-seq2 analysis. C) Comparisons of significantly changed genes from DEseq2 analysis of each mutant compared to wild type (WT) and their *P*-values shown above the arrows. Solid lines and bold text indicate that the overlap is significant (p<0.05), dotted lines and blue text indicate a non-significant *P*-value (p≥0.05).

To assess whether Liv3, Nrg1, and Cqs2 were required for the expression of genes involved in the Qsp1 response, we performed RNA-seq analysis on RNA extracted from wild type, *qsp1Δ, liv3Δ, cqs2Δ,* and *nrg1*Δ cultures grown to OD1 or OD5 in minimal media. We compared differentially expressed genes from the *qsp1*Δ mutant and the three transcription factor deletion strains relative to wild type at both timepoints (Figure 2C). There were significant overlaps between groups of differentially expressed genes under at least one condition (Figure 2C). While the Qsp1-dependent gene set consistently overlapped with those dependent on Cqs2 or Nrg1, this was not the case for the Liv3-dependent set (Figure 2C). The latter only significantly overlapped the Qsp1-dependent set for genes derepressed in the mutants at OD1 (Figure 2C). These data reveal strong similarities between the transcript signatures of *qsp1*Δ mutant and those of *cqs2*Δ and *nrg1*Δ mutants, with only weak similarity to the *liv3*Δ mutant signature.

To further examine the involvement of these three transcription factors in Qsp1 signaling, each transcription factor was tagged with a FLAG epitope tag and expressed from their native promoters in either a wild type or *qsp1*Δ mutant background. Nrg1 was tagged on its N-terminus, as a C-terminal tag rendered the *qsp1*Δ knockout unable to respond to synthetic Qsp1 peptide in the cell wall stress assay.

Quantitative immunoblots show a slight reduction in Cqs2 levels between wild type cells and cells lacking the *QSP1* gene at OD1 in minimal media, though this difference was not significant (Figure 3A and Supplemental Table 1). However, cells lacking *QSP1* expressed significantly more Nrg1 and Liv3 protein (Figure 3A and Supplemental Table 1). Due to the strong overlaps between genes controlled by Qsp1 and these three factors at OD1 in minimal media, we conducted ChIP-seq under this condition (Figure 3B, 3C). To quantify transcription factor binding, a ChIP score for each gene was calculated as the sum of the read depth over a 1 kb region upstream of each transcription start site, normalized to the untagged strain. We employed a k-means clustering approach to divide the genes into groups whose promoters were significantly bound or not-bound (See Methods). We found that in wild type, these transcription factors generally bind to the same promoters (Figure 3C, 3D and Supplemental Table 2). The majority of the genes bound by any transcription factor are also bound by one or two others, with 274 genes bound by all three transcription factors (Figure 3D). The overlaps between the sets of promoters that are bound by any two of these three transcription factors are highly significant, further supporting the conclusion that these transcription factors are part of a network (Figure 3E and Supplemental Table 2).

**Figure 3.**
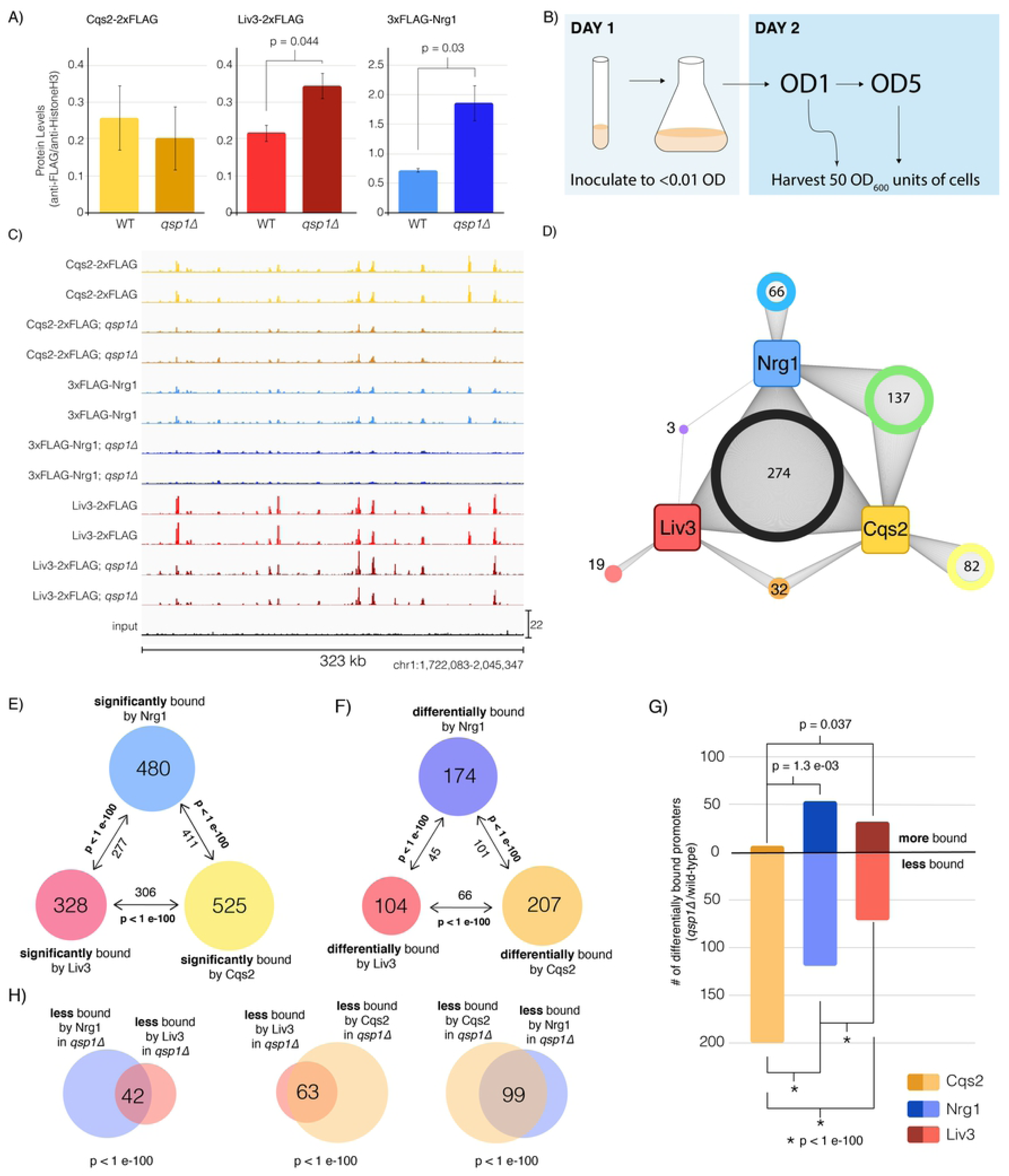
Qsp1 affects Cqs2, Liv3, and Nrg1 binding to a common set of promoters. A) Expression of Cqs2, Liv3, and Nrg1 protein levels in wild type or *qsp1*Δ mutants at OD1 in minimal media. Two biological replicates are shown for each condition. B) Schematic for how ChIP-seq cultures were grown and harvested. C) ChIP-Seq data was visualized using the Integrative Genomics Viewer software. Binding across part of chromosome 1 is shown. D) Network diagram of promoters bound by Nrg1, Liv3, and Cqs2 in wild-type cells in YNB at OD1. E) Overlaps between promoter sets significantly enriched for Liv3, Cqs2, or Nrg1 binding and their significance. F) Overlaps between promoter sets differentially bound (>1.5-fold) by Liv3, Cqs2, or Nrg1 in the *qsp1*Δ mutant compared to wild-type. G) Breakdown of promoters that are bound more or less by a given transcription factor in *qsp1*Δ compared to wild type, filtered by promoters significantly bound in either genotype. Significant overlaps between groups are noted with the *P*-value. H) Overlaps between promoter sets that are >1.5-fold less bound by Liv3, Cqs2, or Nrg1 in a *qsp1*Δ mutant compared to wild type, and their significance. Only promoters that are called as bound in either genotype by k-means analysis are shown.

### Qsp1 affects the binding of all three transcription factors to a common set of promoters

To test whether Qsp1 could influence the binding of these transcription factors, we constructed tagged transcription factor strains that also harbor a knockout of *QSP1*, and conducted ChIP-seq. In specific regions, binding of the transcription factors is abolished or diminished in the absence of *QSP1,* indicating that Qsp1 peptide is required for binding of these factors to these promoters (Figure 3C). Our analysis revealed that Qsp1 affects the binding of Cqs2, Liv3, and Nrg1 to a large fraction of bound promoters (Figure 3F, Supplemental Figure 1 and Supplemental Table 1). We confirmed this by quantifying the amount of binding of each factor to two of the promoters that showed the highest differences by ChIP-seq using ChIP-qPCR (Supplemental Figure 2). Overall, we observed a shift to lower levels of binding in the *qsp1*Δ mutant by Cqs2, whereas Nrg1 and Liv3 binding increased for some promoters and decreased for others (Figure 3G, Supplemental Figure 1, and Supplemental Table 2). In the majority of cases, Qsp1 promotes rather than inhibits the binding of a transcription factor to their targets. This indicates that Qsp1 promotes the binding of Cqs2 upstream of genes and affects the binding of Nrg1 and Liv3.

We next sought to understand whether Qsp1 influences the binding of Cqs2, Nrg1, and Liv3 to the same sets of promoters, and whether this influence was positive or negative. We examined the degree of overlap between the sets of promoters that displayed a mean fold-change greater than 1.5-fold in either direction that were called as bound in either wild-type or *qsp1*Δ mutants and tested whether the overlaps between these sets were significant. We observed highly significant (*P* < 1×10^-100^) overlaps between promoters that displayed lower levels of binding by any two transcription factors in the *qsp1*Δ mutant, more so than between groups of promoters that are more bound in the *qsp1*Δ mutant (Figure 3G, 3H, and Supplemental Table 2). This indicates that Qsp1 functions to promote the binding of all three transcription factors upstream of a subset of genes. The overlaps between genes that are differentially bound by any two of these three transcription factors in the *qsp1*Δ mutant are also highly significant (Figure 3F and Supplemental Table 2), but most of the significance comes from genes that are less bound in the mutant (Figure 3G and Supplemental Table 2).

Cqs2, Nrg1, and Liv3 are transcription factors that bind to DNA and influence gene expression, therefore we tested whether the binding of each transcription factor upstream of a gene impacts the expression of that gene. We compared genes bound by a tagged transcription factor in wild type to genes whose expression was affected by loss of the corresponding transcription factor. The overlaps between these two sets were significant in all comparisons, supporting the conclusion that these three transcription factors influence gene expression via binding to target genes under the conditions tested (Figure 4A and Supplemental Table 1). Non-overlapping genes may be regulated indirectly or via other inputs.

**Figure 4.**
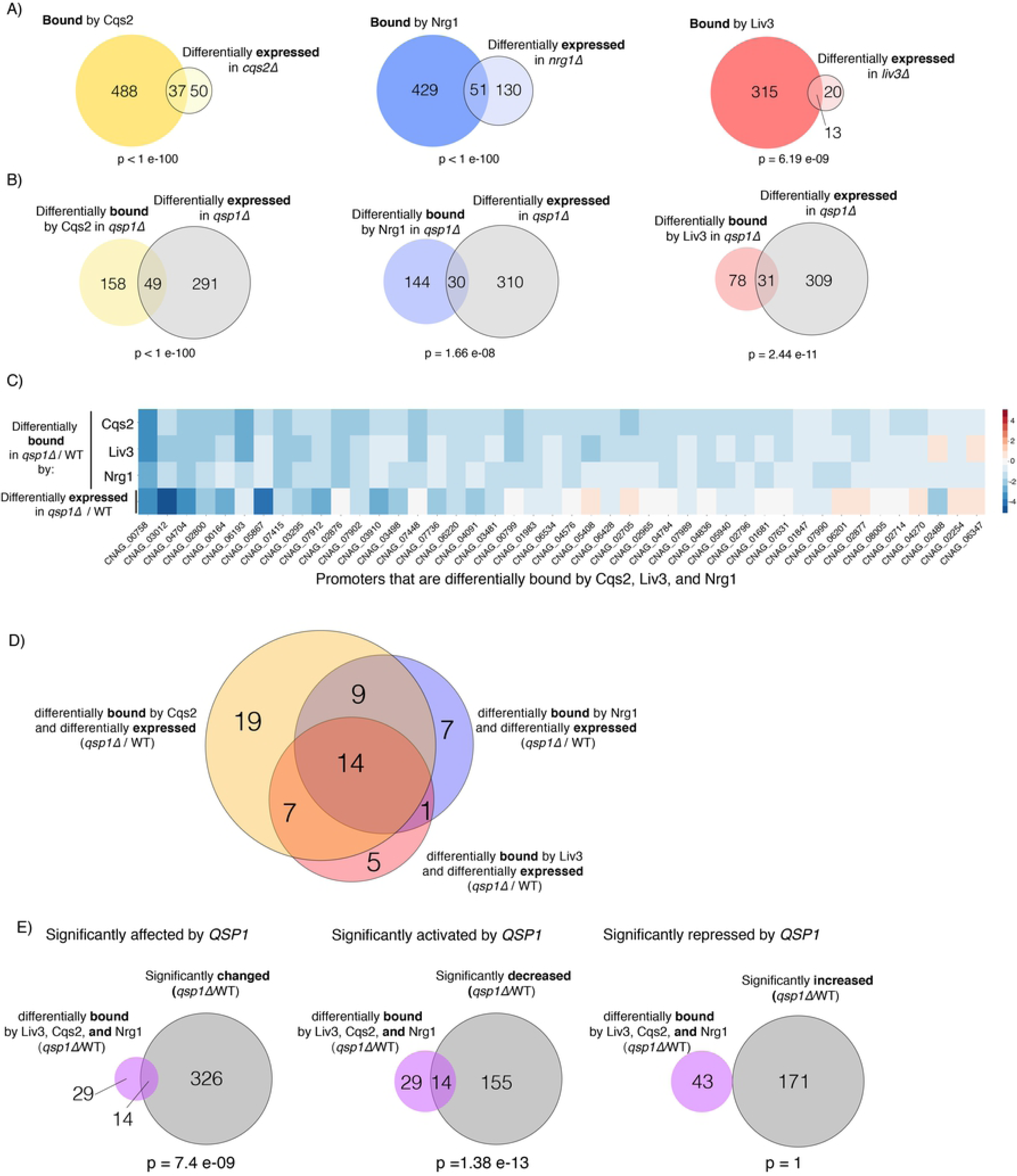
Qsp1 regulation of binding of Cqs2, Nrg1, and Liv3 to promoters correlates with a change in expression. A) Overlap between promoters bound by each transcription factor in wild typeand genes that are differentially expressed in the corresponding transcription factor mutant, at OD1 in minimal media. B) Overlap between promoters differentially bound by each transcription factor and genes that are differentially expressed in each *qsp1*Δ mutant relative to wild type, at OD1 in minimal media. C) Heatmap of genes that are differentially bound by all three TFs Cqs2, Liv3, or Nrg1, and their respective log2-fold expression difference in *qsp1*Δ compared to wild type. Non-significant differences are colored in white, significant decreases in mutant are shown in blue, and significant increases in *qsp1*Δ over wild type are shown in yellow. D) Degree of overlap between sets of genes that are differentially bound by a transcription factor and genes that are differentially expressed in a *qsp1*Δ mutant relative to wild-type, and those that are differentially bound by another transcription factor and differentially expressed in *qsp1*Δ mutants. E) Overlap between gene sets that are differentially bound by all three transcription factors (Liv3, Cqs2, and Nrg1) in the *qsp1*Δ mutant compared to wild type and genes that are significantly changed, decreased, or increased in the *qsp1*Δ mutant over wild type.

### Qsp1 promotes the binding of Cqs2, Nrg1, and Liv3 to promoters, which activates expression of these genes

To test whether the Qsp1-dependent binding events had functional consequences, we next investigated whether the genes whose promoters that were differentially bound in a *qsp1*Δ mutant by each transcription factor were also differentially expressed in a *qsp1*Δ mutant under a particular condition for all three transcription factors. For all three transcription factors, we observed significant overlaps between genes differentially bound by a transcription factor and genes that were differentially expressed in a *qs p1*Δ mutant compared to wild type, indicating that the influence of Qsp1 on binding of Cqs2, Nrg1, and Liv3 is important for gene expression (Figure 4B and Supplemental Table 2).

To test if there was a relationship between the combination of transcription factors bound and gene expression, we created a heatmap displaying data for genes whose promoters are differentially bound in a *qsp1*Δ mutant by any transcription factor at OD1, with their corresponding change in expression in a *qsp1*Δ mutant compared to wild type (Supplemental Figure 3 and Supplemental Table 4). We observed that the largest impact on gene expression occurs for genes whose promoters were much less occupied by all three transcription factors together in the *qsp1*Δ mutant (Figure 4C, Supplemental Figure 3 and Supplemental Table 2). This decrease in occupancy of all three transcription factors in a *qsp1*Δ mutant compared to wild type corresponds to a significant decrease in expression of about a third of these genes (Figure 4C, 4D, 4E and Supplemental Tables 2 and 3).

From our conservative analysis, there are fourteen genes where Qsp1 promotion of Cqs2, Nrg1, and Liv3 binding correlates with a significant change in expression in the *qsp1*Δ mutant compared to wild type (Table 1). Five encode predicted transporters of sugars, amino acids, or other types of nutrients. Interestingly, one of the genes encodes Ral2, which is essential for mating in *Schizosaccharomyces pombe* (15). Ral2 activates Ras1, a GTPase that is also activated by Ste6, the alpha mating factor transporter and exchange factor for Ras1 (15, 16). Another of these genes encodes Agn1, a putative α-glucanase, and could be related to the cell wall phenotype of *qsp1*Δ and *cqs2*Δ mutants (17).

**Table 1.**
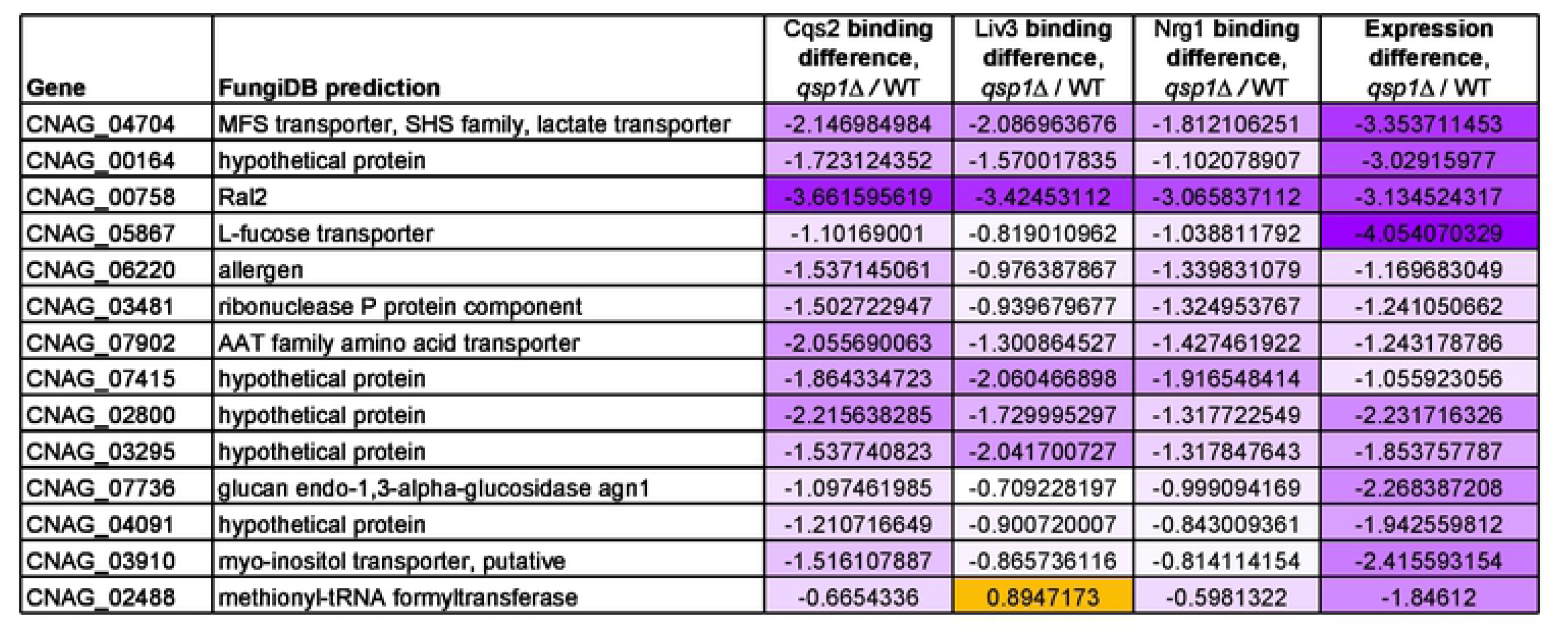
Thirteen genes are differentially bound by Cqs2, Liv3, and Nrg1 and differentially expressed in minimal media at OD1. Predictions are based on the FungiDB database (www.fungidb.org/fungidb).

Together, these data indicate that Qsp1 regulates the binding of Cqs2, Nrg1, and Liv3 together to a subset of Qsp1-regulated genes, and the loss of binding of all three of these factors results in altered expression of a set of genes predicted to be involved in nutrient sensing, signaling, and acquisition as well as cell wall remodeling.

### Loss *NRG1* or *CQS2* affects Liv3, Nrg1, and Cqs2 binding to promoters

To test whether Cqs2, Nrg1, and Liv3 impacted each other’s binding, we attempted to delete the genes encoding the other two transcription factors in the tagged strains described above. We conducted ChIP-seq on FLAG-tagged Liv3, Nrg1, or Cqs2 strains harboring deletions of *NRG1* or *CQS2* (we were unable to obtain deletions of *LIV3*) grown to OD1 in minimal media. Immunoblotting demonstrated that no significant difference in the levels of each of these tagged transcription factors in the mutant background compared to wild type (Figure 5A and Supplemental Table 5). An example of the binding pattern of Liv3, Nrg1, and Cqs2 in these backgrounds across part of chromosome 1 is shown (Figure 5B). We calculated a ChIP score for transcription factor binding for each promoter in each strain and plotted these for each gene in each transcription factor mutant versus wild type (Supplemental Figure 4 and Supplemental Table 6). Strikingly, deletion of *CQS2* results in reduced Liv3 binding to promoters. 357 genes exhibited a >1.5-fold decrease, and only 3 genes exhibited a >1.5-fold increase in binding (Figure 5C, Supplemental Figure 4, Supplemental Tables 2 and 6). In contrast, deletion of *CQS2* both reduced and increased Nrg1 binding, depending on the promoter. Deletion of *NRG1* also dramatically impacted Liv3 binding to targets, again primarily reducing binding. Finally, deletion of *NRG1* increased Cqs2 binding to more targets than it decreased.

**Figure 5.**
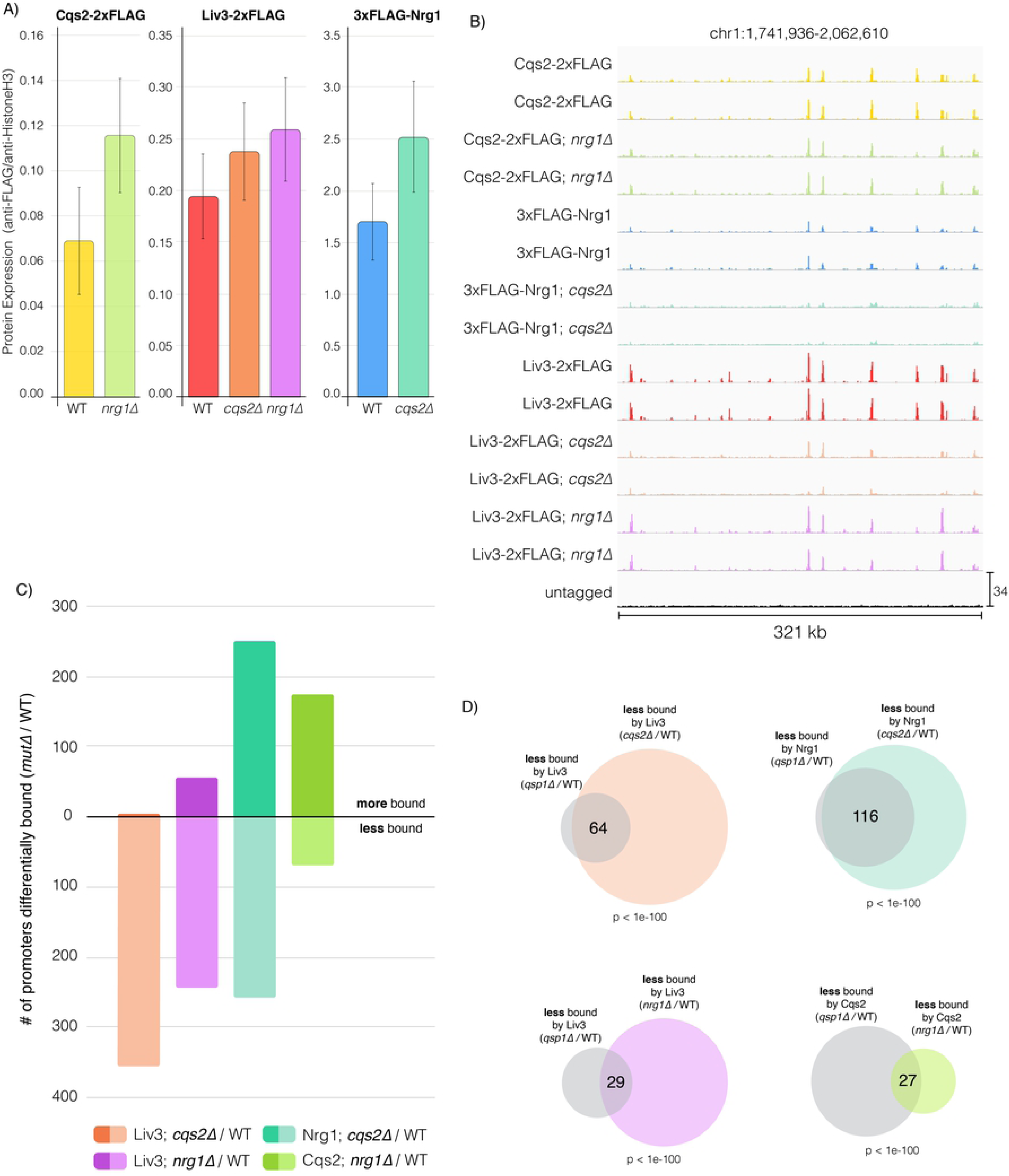
Cqs2 and Nrg1 impact binding of other transcription factors in the network to promoters, and this impact on binding is similar to the impact of Qsp1 on binding. A) Protein levels of FLAG-tagged Cqs2, Liv3, and Nrg1 in each transcription factor deletion strain background. Average of two biological replicates is shown. B) ChIP-Seq data was visualized using the Integrative Genomics Viewer software. Binding across part of chromosome 1 is shown. C) Breakdown of promoters that are bound more or less by a given transcription factor in *qsp1*Δ compared to wild type, filtered by promoters significantly bound in either genotype. D) Overlaps between promoter sets that are less bound by each transcription factor in the *qsp1*Δ mutant and promoter sets that are less bound by each transcription factor in the *nrg1*Δ or *cqs2*Δ mutants.

We sought to understand further the transcription factor network in the context of Qsp1 signaling. To accomplish this, we examined how the loss of *NRG1* or *CQS2* compared with loss of *QSP1* on altering binding of Cqs2, Liv3, and Nrg1 to promoters, by comparing the sets of promoters that were affected by each deletion. Strikingly, almost all of the Qsp1-dependent promoters also exhibit a Cqs2-dependence for binding of Nrg1 and Liv3, and in the same direction (Figure 5D, Supplemental Figure 5, and Supplemental Tables 2 and 6). In other words, the same group of promoters that exhibit altered Nrg1 and Liv3 binding in a *qsp1*Δ knockout is also impacted in the same direction in a *cqs2*Δ knockout. Liv3 and Cqs2 binding to Qsp1-dependent promoters is also significantly regulated by Nrg1, but to a lesser extent (Figure 5D, Supplemental Figure 5, and Supplemental Tables 2 and 6). Thus, there is a notable overlap between promoters whose transcription factor binding is promoted by Qsp1 and those that display transcription factor interdependencies for binding.

## DISCUSSION

Single celled organisms often cooperate in a type of community-oriented signaling called quorum sensing, mediated by the accumulation of secreted autoregulatory molecules. Quorum sensing coordinates cellular adaptations that allow the cells to survive in response to environmental cues, such as in mating, biofilm formation, and host infection. Quorum sensing has been reported in many different microbes to regulate mating and competence (18–22), starvation and mating (23), regulation of sporulation in response to starvation (24), nutrient acquisition and virulence (25). Our experiments have uncovered that Qsp1 regulates gene expression by influencing three transcription factors that play roles in mating, virulence, and nutrient acquisition, providing insight into the mechanism by which a eukaryotic quorum sensing molecule can influence gene expression and clues to the role that quorum sensing plays in the biology of *C. neoformans*.

Our previous work in *C. neoformans* demonstrated that Qsp1 is secreted as a precursor that is cleaved outside the cell (4). The mature peptide accumulates in the culture supernatant, then appears to be imported back into the cell by Opt1, where it induces a transcriptional response (4). This is remarkably similar to some gram-positive bacteria, which also secrete quorum sensing peptides that are imported into the cell via oligopeptide permeases. Once imported, these small peptides interact with phosphatases (26, 27) or transcriptional regulators (28–30) to influence gene expression. In *C. neoformans*, the two transcription factors Liv3 and Cqs2 have been identified as regulators of the Qsp1 response (4, 12). In this study, we uncovered Nrg1 as a third Qsp1-regulated transcription factor. We showed that the response to Qsp1 signaling is mediated by a network formed by these three transcription factors (Figure 2), which were identified on the basis of a temperature-regulated rough colony morphology that is also exhibited by a *qsp1*Δ knockout (Figure 1). Surprisingly, we found that Qsp1 seems to promote the binding of Cqs2 to promoters and alter the binding of Nrg1 and Liv3 (Figure 3). These Qsp1-dependent promoters are shared between all three transcription factors, with the largest and most significant overlaps occurring between promoters to which Qsp1 promotes transcription factor binding (Figure 3). This decrease in transcription factor binding to Qsp1-dependent promoters correlates with a decrease in gene expression in cells lacking *QSP1* compared to wild type (Figure 4). We observed this correlation with reduced binding leading to reduced expression in spite of higher levels of Cqs2 and Nrg1 protein in *qsp1*Δ mutants in this condition (Figure 3). Furthermore, Cqs2 promotes Liv3 binding to promoters (Figure 5). Additionally, Cqs2 impacts Nrg1 and Liv3 binding on almost all Qsp1-dependent promoters, given the strong overlap between promoters affected for binding of these transcription factors by loss of *QSP1* and loss of *CQS2* (Figure 5). Nrg1 also impacts Cqs2 and Liv3 binding (Figure 5). Qsp1 may influence on Nrg1 and Liv3 binding by promoting Cqs2’s affinity for promoter sites, since Cqs2 appears to be the principal factor through which Qsp1 acts under these conditions (Figure 6). Together, these data support the model that Qsp1 influences gene expression by controlling a network of transcription factors’ ability to bind to DNA, (Figure 6). It is unclear if Qsp1 directly binds to Cqs2, or if there is another unidentified Qsp1-regulated signaling factor upstream of Cqs2 that regulates the affinity of Cqs2 for target promoters.

**Figure 6.**
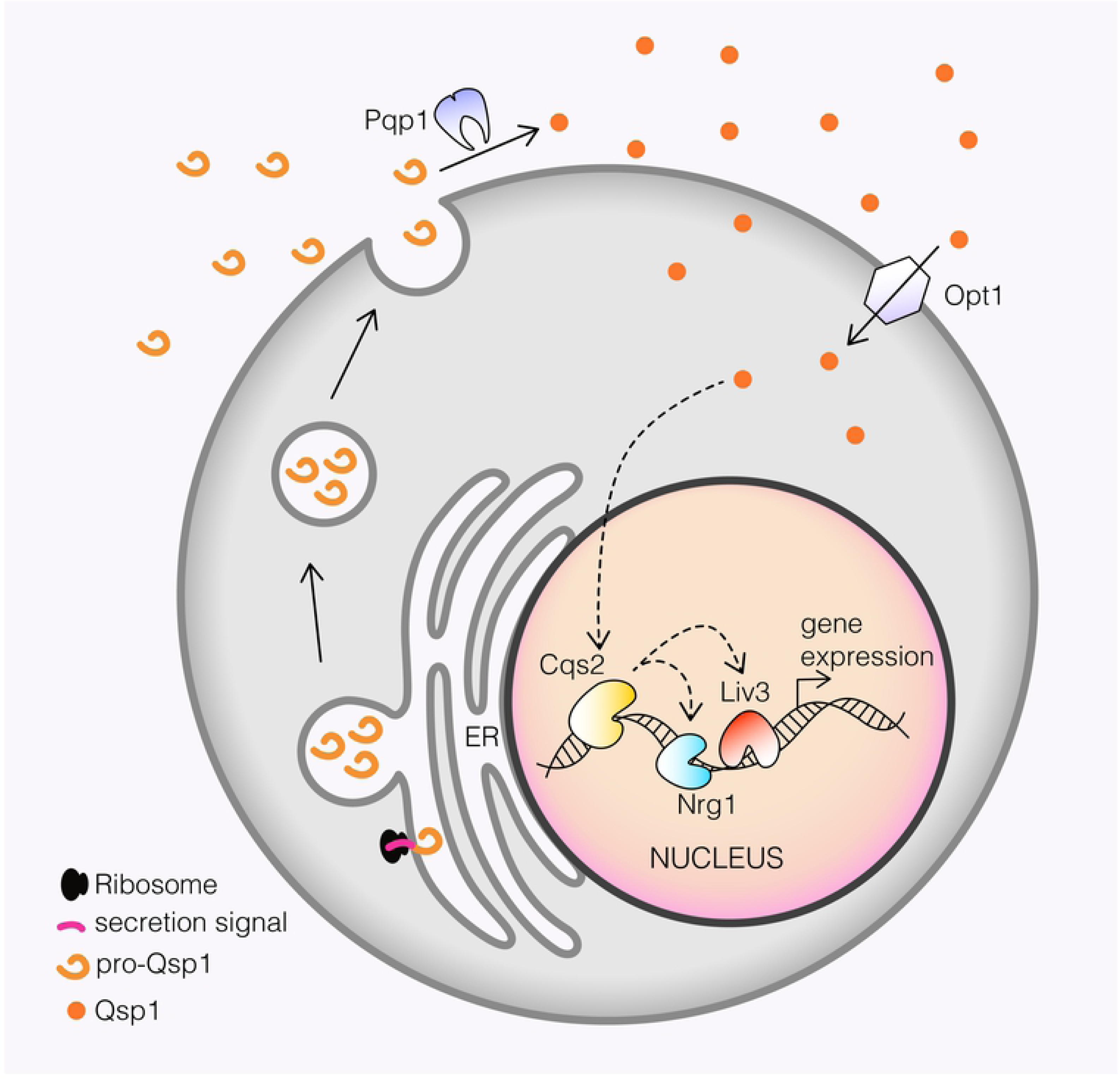
Model for how Qsp1 triggers changes in gene expression in *Cryptococcus neoformans*. Following import into the cytoplasm, Qsp1 alters the binding of Nrg1 and Liv3 by modulating the ability of Cqs2 to bind promoters, thereby causing changes to gene expression. Dotted lines indicate functional rather than physical interactions.

Cqs2, Nrg1, and Liv3 are transcription factors that play roles in mating, nutrient acquisition and virulence in *C. neoformans.* Qsp1 has recently been shown to signal through Cqs2 to regulate unisexual reproduction and filamentation (12). Liv3 is required for proliferation in the lung (11). Liv3 is also a homolog of Wor1, the master regulator of white-opaque switching in *C. albicans*, a functional and morphological switch in phenotype that can be triggered by various environmental cues and determines which area of the body the fungus is best equipped to colonize (6,8,9). Nrg1 is a transcriptional regulator that promotes bisexual mating and virulence, and plays a role in several cellular processes, including carbohydrate acquisition, metabolism, and capsule formation (14). Homologs of Nrg1 in other fungi play roles in filamentation, nutrient sensing, and metabolism in response to environmental cues and are also regulated by quorum sensing (31–37). Here, we show that Nrg1 and Liv3 protein levels are repressed by the *QSP1* gene in minimal media (Figure 3A). We also found that Qsp1 promotes the binding of Cqs2 to promoters and influences the binding of Nrg1 and Liv3, and that these transcription factors influence each other’s binding (Figure 3). These experiments provide a mechanistic basis for quorum sensing control of these factors and further evidence for the implication of quorum sensing in mating and pathogenesis of *C. neoformans*.

It is unclear why Cqs2, Nrg1, and Liv3 transcription factors have been integrated into a quorum sensing system. One possibility is that quorum sensing enables cells to anticipate and prepare for future starvation and associated stresses, which could be critical in particular host niches or when deciding to mate. In prokaryotes, starvation and quorum sensing signaling pathways regulate each other (38). In *Saccharomyces cerevisiae,* the production of autoregulatory aromatic alcohols is coupled to both culture density and nitrogen starvation, and serves as a species-specific trigger for transformation into a filamentous form (39). In both bacteria and yeast, it is thought that entry into stationary phase once a nutrients are exhausted provide benefits to the cell such as thickening of the cell wall, accumulation of reserve nutrients, and an increased resistance to environmental stressors, allowing the cells to survive long term (38, 40). Integration of quorum sensing and starvation signaling could explain why Qsp1 signaling increases as culture density increases in rich media as nutrients run out, but has the opposite trend in minimal media, where cells are starved immediately (Figure 2). In addition, we found that Qsp1 promotion of resistance of stationary phase cells to cell wall stress requires Nrg1 and Cqs2 (Figure 1), further solidifying the relationship between quorum sensing and starvation responses.

In line with this idea, one of the promoters that exhibited a very dramatic dependence on Qsp1 for binding of all three transcription factors was the *LAC1* gene (CNAG_03465) (Supplemental Figure 2), which encodes the melanization factor laccase (41–43). Melanization is known to be a key virulence trait for *C. neoformans* infection (41, 44). In our previous work, we found that cells lacking Qsp1 display altered capacities to produce melanin when plated on plates containing the substrate for melanin production (4). Although our conditions did not reveal laccase transcript regulation, it may be that these three transcription factors bind to the laccase promoter in the presence of Qsp1 in order to prime the cell for transcription when additional signals are received.

In conclusion, it seems that these three transcription factors are at the core of a gene regulatory network that integrates Qsp1 signaling with starvation or other unknown signaling inputs to decide which genes to express in different contexts (such as in the host or in different media), ultimately influencing the mating and virulence of this organism.

## METHODS

### Cryptococcal strain construction

Gene deletions were generated using nourseothricin (NAT) resistance, neomycin (NEO) resistance, or hygromycin (HYG) resistance cassettes. Proteins were tagged with 2x-FLAG or 3x-FLAG epitope tags using one of these three resistance cassettes as previously described (44). Constructs were made via homologous recombination using fragments amplified with the primers in Table 2. Strains constructed in this study are listed in Table 3. All strains are derived from the KN99alpha (CM26) parent.

**Table 2.**
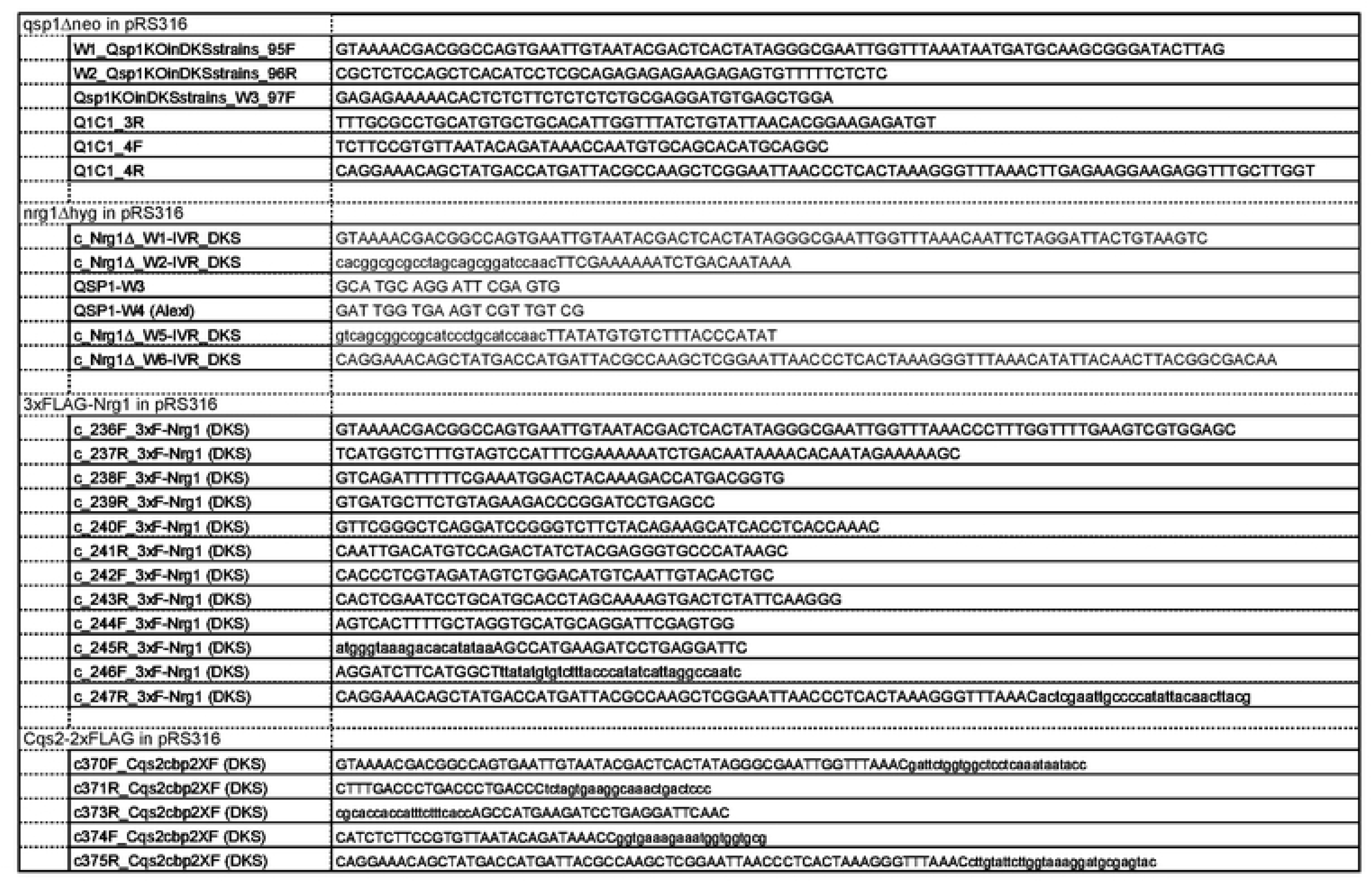
Primers used to create genetic constructs to create strains used in this study

**Table 3.**
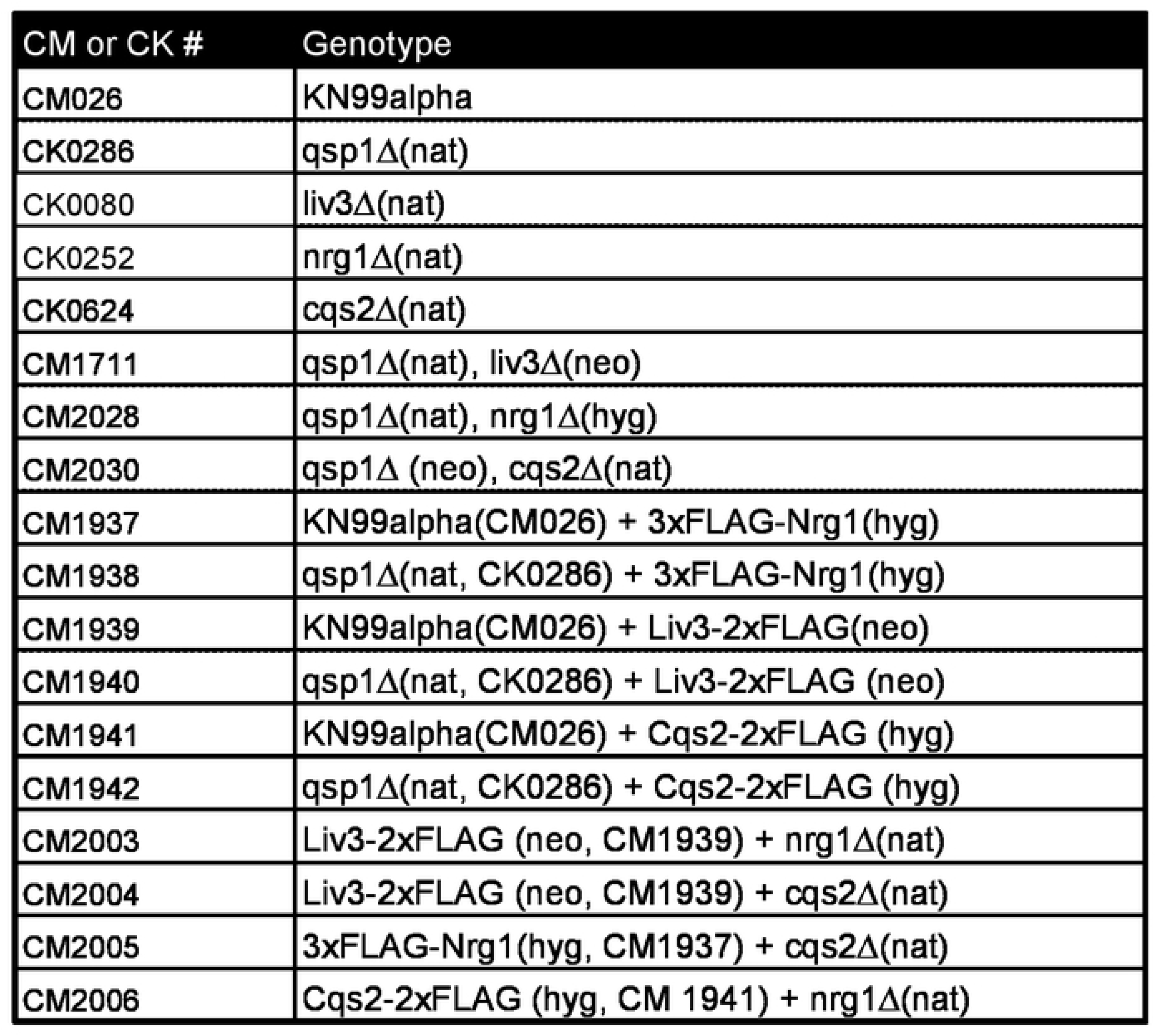
Strains used in this study.

### Cell Wall Stress Assay

Part of a colony was cultured in 5 ml of YPAD (1% yeast extract, 2% Bacto-peptone, 2% glucose, 0.015% L-tryptophan, 0.004% adenine) for 48 hours, when all cultures were fully saturated. 1 uM pure synthetic Qsp1 peptide (LifeTein) was added to indicated cultures at the time of inoculation. 50 ul of saturated culture was mixed with 150ul of either water or SDS in a series of 1:2 dilutions (starting at 10%) and incubated for 3 hours at room temperature without shaking. The supernatant of settled incubations was partially replaced with water following SDS incubation and prior to cell resuspension to minimize the amount of SDS transferred to the recipient plate, and 3 ul were spotted on YPAD plates containing no SDS to assay survival. Cells incubated in water were then serially diluted 1:6 to provide a measure of the titer of the input culture.

### Immunoblots

Cultures were grown as indicated. 2 OD_600_ units of cells per sample were fixed with 10% TCA, then 100% acetone, then lysed by two 1.5min rounds of bead-beating in sample buffer. Samples were then boiled for 5 min and cell debris was spun down. For supernatant analysis, 2 mL of conditioned media were snap frozen and lyophilized overnight, then resuspended in 150 ul of 1x Laemmli Sample Buffer. 5-10 ul of each sample was loaded on 4-12% Bis-Tris gels (Thermofisher).

### RNA-Seq and ChIP-Seq cultures

Each strain of C. neoformans was inoculated in YPAD or YNB at 30°C. The next day, larger cultures were started from the starter cultures at an OD < 0.01. On the following day, as each culture grew in density 50 OD_600_ units of cells were harvested sequentially from the same culture at the indicated ODs.

50 OD_600_’s of cells at each optical density (OD 1, 5, and 10) for each replicate for each strain were harvested sequentially as the cultures grew. For ChIP-seq samples, 50 OD_600_’s of cells were crosslinked in a 50 mL volume of conditioned media from the same culture, harvested at the same time as the cells.

### RNA-Seq

Total RNA was isolated from 50 OD_600_’s of cells as previously described (45) and libraries prepared as previously described (46). In brief, cell pellets were lyophilized overnight and then RNA was isolated using TRIzol (Invitrogen) as previously described (45) and DNase treated as previously described (47). 0.5 ug RNA was then prepared for sequencing using the QuantSeq 3‘-mRNA-Seq Library Prep Kit FWD (Lexogen) according to the manufacturer’s instructions. Input RNA quality and mRNA purity were verified by Bioanalyzer Pico RNA chips (Agilent). Libraries were sequenced on the HiSeq 4000 platform (Illumina).

### RNA-Seq Analysis

Expression analysis for each transcription factor mutant was performed by counting the number of reads aligned by STAR for each transcript (48). DEseq2 was used to determine genes differentially expressed between mutant and wild type conditions. See GEO accession #GSE147378 for raw data.

### Chromatin immunoprecipitation (ChIP)

ChIP was performed as described previously (4), with the following changes: 50 OD_600_ units of cells were crosslinked in 50 mL total of conditioned media. Lyophilized pellets were resuspended in 600 ul ChIP lysis buffer with protease inhibitors for bead beating until >95% of cells were lysed. The chromatin pellet was resuspended in 350 ul ChIP lysis buffer for sonication. After sonication and removal of cell debris via centrifugation, the supernatant was brought to 3 ml in ChIP lysis buffer. Immunoprecipitation was performed at 4°C overnight with nutation in 1 ml chromatin aliquots with 3 ul of anti-FLAG M2 antibody (F3165, Sigma) and 20 ul of Protein G Dynabeads (Invitrogen). See GEO accession #GSE147378 for raw data.

### ChIP-seq library construction

ChIP-seq library construction was performed as described previously (4) with the following changes: For each genotype, libraries for two biological replicates were prepared. Adaptors were selected out using Sera-Mag SpeedBead Carboxylate-Modified Magnetic Particles (Hydrophobic) and products between 200-500bp were selected by gel extraction. Library quality and concentration were determined by High Sensitivity DNA Bioanalyzer analysis (Agilent) and Qubit (Thermofisher), respectively.

### ChIP-Seq analysis

ChIP-seq reads were trimmed with CutAdapt and aligned using Bowtie1 (49). Up to two mismatches within the seed sequence. If a read could align to multiple loci, it was aligned at random. Indexed, sorted bam files were created for each dataset using SAMtools (50). Bedgraph files were created using BEDtools (51, 52), and normalized to the untagged sample by subtraction. The ChIP signal for each gene in each replicate was calculated as the sum of the read depth over the promoter region, defined as the 1kb upstream of the annotated transcription start site. Using the ChIP signal, genes were clustered into bound or unbound via k-means analysis. There was a high degree of overlap between genes called as bound in either replicate (Supplemental Table 7), but genes were only considered as bound in our analysis if it was called as bound in both replicates by k-means analysis (Supplemental Table 6). The average of the ChIP signal from both replicates was taken for subsequent analysis. Bedgraphs were plotted using Integrative Genomics Viewer 2.0.30 (Broad Institute). Genes were determined as differentially bound if the fold change in ChIP signal between both conditions was greater than 1.5-fold in either direction (based on (53)).

### Statistical Analysis

Immunoblot quantification was performed with ImageJ analysis, and significance was determined using the student’s T-test. *P-*values <0.05 were considered significant. All other *P-*values were adjusted (P-adj) for multiple testing using the Bonferroni correction (Supplementary Table2).

## ACKNOWLEDGEMENTS

We thank members of the Madhani lab for helpful discussions and review of the paper, especially Sandra Catania, Jordan Burke, and Daniele Canzio. We thank Nguyen Nguyen for media preparation. This work was supported by R56AI126726 and R01AI20464. DKS and HDM designed the study. DKS and BR performed the experiments, DKS and DSP performed the bioinformatics analysis, and HDM supervised the work.

## SUPPLEMENTAL FIGURE LEGENDS

Supplemental Figure 1. Deletion of *QSP1* affects the binding of Cqs2, Liv3, and Nrg1 to promoters.

ChIP score for each gene was calculated as the read depth in the 1 kb region upstream of the transcription start site, normalized to the untagged control. Only promoters that are called as bound in either genotype by k-means analysis are shown, with promoters that are more or less bound (>1.5-fold changed) by each factor in the *qsp1*Δ mutant highlighted in orange or light blue, respectively. The number of promoters in either of these groups is labeled with the corresponding color.

Supplemental Figure 2. ChIP-qPCR validation of ChIP-seq results.

ChIP was performed on tagged strains followed by qPCR to quantify binding of CNAG_00758 and CNAG_03465 by tagged Nrg1, Liv3, and Cqs2 in wild type or *qsp1*Δ knockout using the primers in Table 4.

**Table 4.**
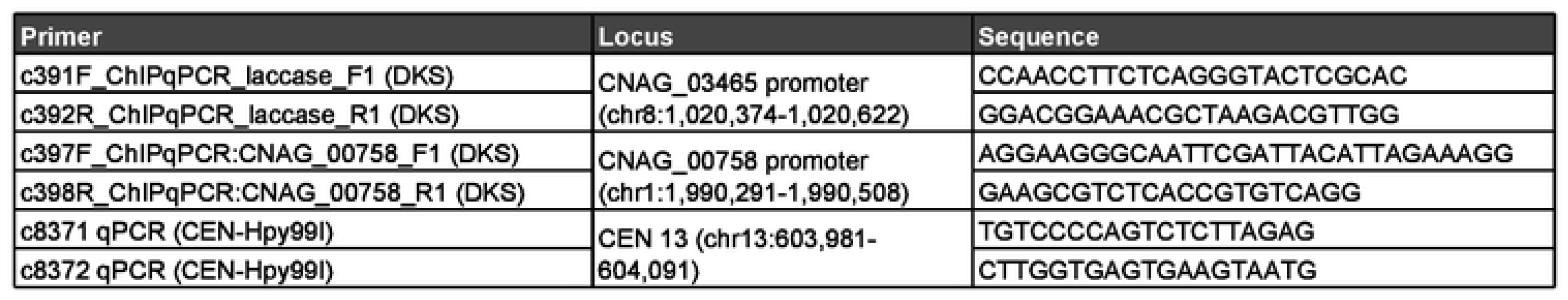
Primers used for ChIP-qPCR.

Supplemental Figure 3. A heatmap of genes that are differentially bound by either Cqs2, Liv3, or Nrg1, and their respective log2-fold expression difference in *qsp1*Δ compared to wild type.

Non-significant differences are colored in white, significant decreases in mutant are shown in blue, and significant increases in *qsp1*Δ over wild type are shown in yellow.

Supplemental Figure 4. C*Q*S2 or *NRG1* deletion affects the binding of Cqs2, Liv3, and Nrg1 to their promoters.

The ChIP score for each gene was calculated as the read depth in the 1 kb region upstream of the transcription start site, normalized to the untagged control. Only genes that are called as bound in either genotype by k-means analysis are shown, with genes that are differentially bound by each factor in mutant compared to wild-type (greater than 1.5-fold changed) highlighted in light blue.

Supplemental Figure 5. Promoters that are differentially bound by transcription factors in a *qsp1*Δ mutant are also differentially bound by these transcription factors in an *nrg1*Δ or *cqs2*Δ mutant compared to wild type.

The ChIP score for each gene was calculated as the read depth in the 1 kb region upstream of the transcription start site, normalized to the untagged control. Only genes that are called as bound in either genotype by k-means analysis are shown, with genes that are differentially bound (>1.5-fold changed) by each transcription factor in the *qsp1*Δ mutant highlighted (light blue and yellow). Promoters that are differentially bound by each transcription factor in the transcription factor mutant are highlighted in yellow.

## SUPPLEMENTAL TABLE LEGENDS

Supplementary Table 1. Data used to generate figure 3A.

Quantification of bands in fluorescent immunoblots was performed using the ImageJ gel analysis tool. A student’s T-test was performed to determine significance of the difference in averages of transcription factor expression between wild type and *qsp1*Δ mutant conditions.

Supplementary Table 2. Analysis of the degree and significance of overlaps between different sets of genes either bound by Nrg1 (N), Liv3 (L), or Cqs2 (C) in each genotype, differentially bound by these three transcription factors in *qsp1*Δ vs. wild type, or differentially expressed in *qsp1*Δ vs. wild type.

The figure generated from the data is indicated. Analysis of differentially bound (diff), decreased in binding level (less bound), or increased in binding levels (more bound) of these transcription factors to sets of genes in different mutants compared to wild type is included.

Supplementary Table 3. Values used to create Supplemental Figure 3.

Set of genes that are differentially bound by either Cqs2, Liv3, or Nrg1, and their respective log2-fold expression difference in qsp1Δ compared to wild type. Non-significant differences are given a value of “NA”, significant decreases in mutant are shown in blue, and significant increases in qsp1Δ over wild type are shown in yellow.

Supplementary Table 4. Numbers used to generate the heatmap in Figure 4C & 4D.

Genes that are differentially bound by all three transcription factors are shown, with their fold change in Cqs2, Nrg1, or Liv3 binding in *qsp1*Δ mutants vs wild type, and the gene’s respective log2fold expression change in *qsp1*Δ mutant vs wild type.

Supplementary Table 5. Data used to generate figure 5A.

Quantification of bands in fluorescent immunoblots was performed using the ImageJ gel analysis tool. A student’s T-test was performed to determine significance of the difference in averages of transcription factor expression between wild type and mutant conditions.

Supplementary Table 6.

Values of ChIP scores for binding of each transcription factor in each genotype for each promoter, whether the promoter was called as bound by k-means analysis (“1”) or not (“0”), values for fold change in binding levels in differential binding analysis, and log2fold-change for expression of the gene downstream of the promoter in *qsp1*Δ vs. wild type by RNA-seq followed by DE-seq analysis.

Supplementary Table 7. Comparison of the number of promoters called as bound in either replicate.

Bound genes in each sample were determined by k-means analysis.

